# Low miRNA abundance disables microRNA pathway in mammalian oocytes

**DOI:** 10.1101/757153

**Authors:** Shubhangini Kataruka, Martin Modrak, Veronika Kinterova, Daniela M. Zeitler, Radek Malik, Jiri Kanka, Gunter Meister, Petr Svoboda

## Abstract

MicroRNAs (miRNAs) are ubiquitous small RNAs, which guide post-transcriptional gene repression in countless biological processes. However, the miRNA pathway in mouse oocytes appears inactive and is dispensable for development. We report that miRNAs do not accumulate like maternal mRNAs during oocyte growth. The most abundant miRNAs total tens of thousands of molecules in fully-grown oocytes, a number similar to that observed in much smaller fibroblasts. The lack of miRNA accumulation acts like miRNA knock-down, where miRNAs can engage their targets but are not abundant enough to produce significant silencing effect. Injection of 100,000 miRNAs was sufficient to restore reporter repression in oocytes, confirming that miRNA inactivity primarily stems from low miRNA abundance and not from an active repression of the miRNA pathway *per se*. Similar situation was observed in rat, hamster, porcine, and bovine oocytes arguing that miRNA inactivity is not a mouse-specific adaptation but a common mammalian oocyte feature.

## INTRODUCTION

MicroRNAs (miRNAs, reviewed in detail in^1^) are genome-encoded small RNAs, which guide post-transcriptional repression of gene expression. Their biogenesis starts with nuclear processing of long primary transcripts from which small hairpin miRNA precursors (pre-miRNAs) are released. Pre-miRNAs are then transported to the cytoplasm and cleaved by RNase III Dicer into 21-23 nucleotide RNA duplexes. One strand of the duplex is then loaded onto an Argonaute (AGO) protein, the key component of the effector complex repressing cognate mRNAs (reviewed in^2^). The functional miRNAs in somatic cells have a relatively high copy number^3, 4, 5^. A HeLa cell contains ~50,000 *let-7* miRNA molecules^3^, which is just an order of magnitude below ~580,000 mRNAs estimated to be present in a HeLa cell^6^.

There are two distinct modes of target repression. RNA interference (RNAi)-like mode requires a miRNA loaded on AGO2 and formation of a perfect or nearly perfect miRNA:mRNA duplex. In this case, exemplified by mammalian mir-196 and HoxB8 mRNA^7^, AGO2 makes endonucleolytic mRNA cleavage in the middle of the miRNA:mRNA duplex. However, mammalian miRNAs loaded on one of the four mammalian AGO proteins (AGO1-4) typically bind their cognate mRNAs with imperfect complementarity. Functional miRNA:mRNA interaction may involve little beyond the “seed” region comprising nucleotides 2 to 8 of the miRNA^8, 9, 10, 11^. This results in translational repression^12, 13^ followed by substantial mRNA degradation^14^, which is typical mammalian miRNA mode of target repression.

Thousands of miRNAs have been annotated in mammals^15^. Experimental data suggest that miRNAs regulate directly or indirectly thousands of genes in one cell type^14, 16, 17, 18^ and that a single miRNA might directly suppress hundreds to over a thousand of mRNAs^15, 16^. It has been estimated that miRNAs may collectively target over 60% of mammalian genes^19^.

Mammalian miRNAs were implicated in the majority of cellular and developmental processes and changes in their expression are observed in various diseases^1^. One of the notable exceptions are mouse oocytes where it was found that miRNAs are present^20, 21^, but are unable to repress their targets and are dispensable for early development^22, 23^.

It was proposed that miRNA inactivity could stem from inefficient formation of the miRNA effector complex^24^ or could be linked to an alternative oocyte-specific AGO2 protein isoform^25^. Since kinetic studies of miRNA-mediated repression in somatic cells suggest that miRNA and target concentrations are critical factors for efficient repression^10, 11^, we hypothesized that stoichiometry could be the key factor contributing to inefficient miRNA-mediated repression in the oocyte, which is one of the largest mouse cells.

While a somatic cell has a diameter of 10-20 μm, a meiotically incompetent mouse oocyte grows during the ~2.5 weeks-long growth phase from a diameter of 40 μm to a 80 μm. Consequently, cytoplasmic volume of a fully-grown germinal vesicle intact (GV) oocyte (~260 pl) is almost two orders of magnitude larger than that of a somatic cell like NIH3T3 fibroblast (denoted 3T3 hereafter), which is ~3 pl (Table 1). During the growth phase, a mouse oocyte accumulates up to ~27 million mRNA molecules while an average somatic cell contains 100,000 to 600,000 mRNAs per cell^6, 26, 27, 28, 29, 30^. Importantly, cytoplasmic mRNA concentration in fully-grown oocytes and somatic cells is similar (100-300 nM, Table 1). Maternal mRNA accumulation during the oocyte growth phase is facilitated by extended average mRNA half-life of ~10 days^31, 32, 33^, presumably reflecting reduced capacity of mRNA degradation.

**Table 1.** Quantitative aspects of mRNAs and miRNAs in somatic cells and oocytes.

|  | NIH 3T3 | growing oocyte | fully-grown oocyte |
| --- | --- | --- | --- |
| diameter [$\mu\text{m}$] | 18<sup>1</sup> | 40 | 80 |
| total volume [pL] | 3 | 33.5 | 270 |
| cytoplasmic vol. [pL] | 2.95 | 23.5 | 260 |
| number of molecules | cytoplasmic molar concentration [nM] |  |  |
| 25,000 | 14 | 1.7 | 0.15 |
| 50,000 | 28 | 3.4 | 0.3 |
| 100,000 | 56 | 6.8 | 0.6 |
| 250,000 | 140 | 17 | 1.5 |
| 500,000 | 280 | 34 | 3.0 |
| 5,000,000 | 2,800 | 340 | 30 |
| 25,000,000 | 14,000 | 1,700 | 150 |

<sup>1</sup> Countess™ NIH/3T3 cell data sheet/B10NUMB3R5, ID #108905

Here we report that miRNAs do not accumulate during oocyte growth, resulting in two orders of magnitude shift in miRNA:mRNA ratio, which is likely the principal cause of apparent inactivity of maternal miRNAs. Maternal miRNAs can apparently engage their targets but are not abundant enough to produce significant silencing impact. The same dilution phenomenon has been observed in four other mammalian oocytes. Finally, miRNA activity can be restored in mouse oocytes by injecting as few as 100,000 miRNA molecules. Altogether, our data support a model where miRNA inactivity primarily stems from low miRNA abundance caused by the lack of miRNA accumulation during oocyte growth.

## RESULTS

### miRNA amount in mouse oocytes does not increase during the growth phase

To assess miRNA abundance in oocytes, we analyzed small RNA sequencing (RNA-seq) data^20, 34, 35^ and selected eight different miRNAs either because of their high abundancy in oocytes or because they were unique, so there were no family members that could interfere with their analysis. Next, we quantified these miRNAs in 3T3 fibroblasts, meiotically incompetent mouse oocytes, and fully-grown GV mouse oocytes by qPCR (Fig. 1A). Particular care was taken of potential biases in producing calibration curves and primer specificity was examined as well (Fig. S1).

**Figure 1.**
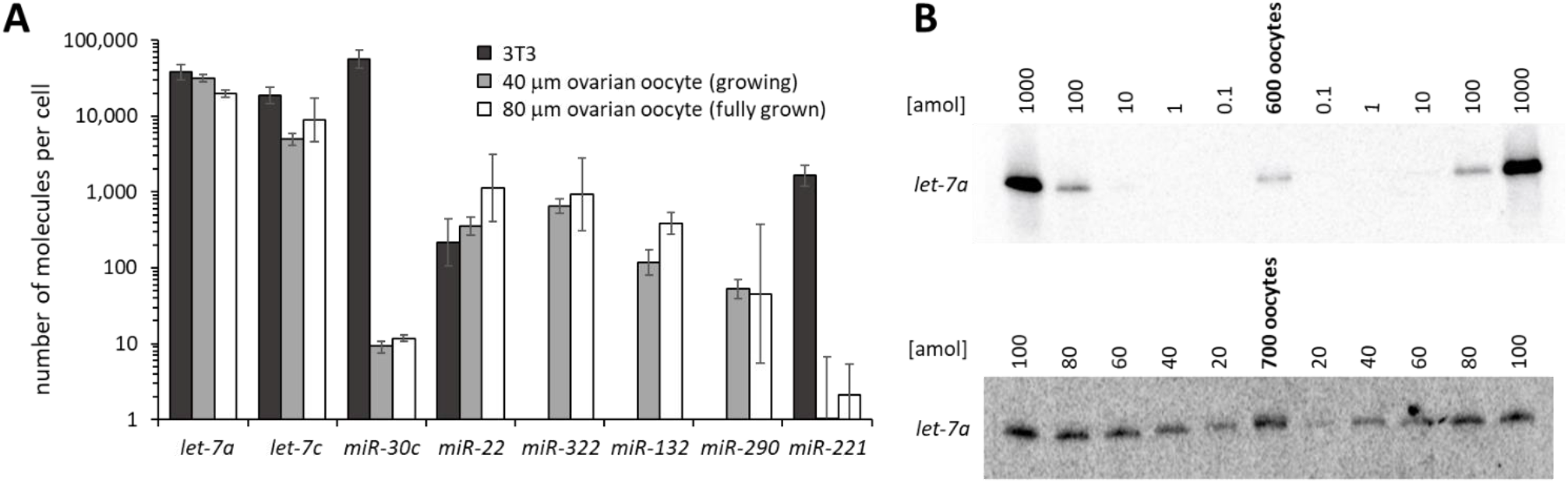
miRNA quantification in oocytes and 3T3 cells. (A) qPCR-based quantification of miRNAs in 3T3 (Ø 18 μm), mouse growing oocyte (Ø 40 μm) and fully-grown GV oocyte (Ø 80 μm). miRNA copy numbers per cell were estimated in three independent experiments using the standard curve in Fig. S1A. Error bars = standard deviation (SD). X-axis represents logarithmic scale. (B) Northern blots for quantifying *let-7a* miRNA counts in mouse fully-grown GV oocytes. Serially-diluted *let-7a* synthetic oligonucleotide was used for building a standard calibration curve (Fig. S1).

Our results revealed that among the analyzed miRNAs in 3T3 cells and oocyte stages the most abundant miRNAs were present at tens of thousands of copies (Fig. 1A and Table S1). This suggests that, in contrast to maternal mRNA accumulation, oocytes do not accumulate miRNAs during the growth phase. This is consistent with similar relative amounts of *let-7* miRNAs observed between P15-16 (growing) and P20-21 (fully-grown) oocytes previously^36^. We estimated by qPCR that *let-7a*, the most abundant of our studied maternal miRNAs, would have ~20,000 copies/oocyte. To validate qPCR data, we analyzed maternal *let-7a* directly by quantitative northern blotting (Fig. 1B), which does not include reverse transcription and geometric amplification of the signal like qPCR. Northern blot analysis estimation yielded ~ 65,600 *let-7a* molecules per oocyte in the absence of a detectable miRNA precursor signal (Fig. 1B). While this number is higher than the qPCR-based estimate, we cannot rule out that other *let-7* family members contributed to the higher estimation. In any case, northern blot also showed that maternal *let-7a* biogenesis is intact and *let-7a* concentration in the oocyte is in sub-nanomolar range.

Taken together, while the concentration of mRNAs in the cytoplasm is comparable for oocytes and 3T3 cells (100-300 nM), similar numbers of miRNA molecules in somatic cells and oocytes make miRNA:mRNA stoichiometry unfavorable in oocytes. For example, 50,000 miRNA molecules in 3T3 cells would be ~28 nM but in mouse oocytes only ~0.3 nM, hundred times less (Table 1).

### Lack of miRNA accumulation is common in mammalian oocytes

An alternative explanation for the lack of accumulation of maternal miRNAs during oocyte growth could be the presence of the endogenous RNA interference (RNAi) pathway, which uses small interfering RNAs (siRNAs) loaded also on AGO proteins. Endogenous RNAi evolved in rodents and is highly active and essential in mouse oocytes^23, 36, 37, 38^. *Ago2*, which is essential for endogenous RNAi in mouse oocytes^39^, has specifically reduced expression level in mouse oocytes and produces a truncated transcript and protein^25^. It could be possible that mouse oocytes represent a unique case where miRNAs do not accumulate because they compete with siRNAs for limited amount of AGO proteins.

To examine this possibility, we quantified maternal miRNAs in four other mammalian species, including bovine and hamster oocytes where the truncated *Ago2* transcript does not exist according to RNA-seq data^40, 41^. Analysis of rat, golden hamster, bovine, and porcine oocytes showed miRNA tallies similar to those in mouse oocytes (Fig. 2, Table 1). While rodent oocytes (Fig. 2A) had somewhat lower miRNA counts than porcine and bovine oocytes (Fig. 2B), even the highest numbers remained around 50,000 miRNA molecules per oocyte.

**Figure 2.**
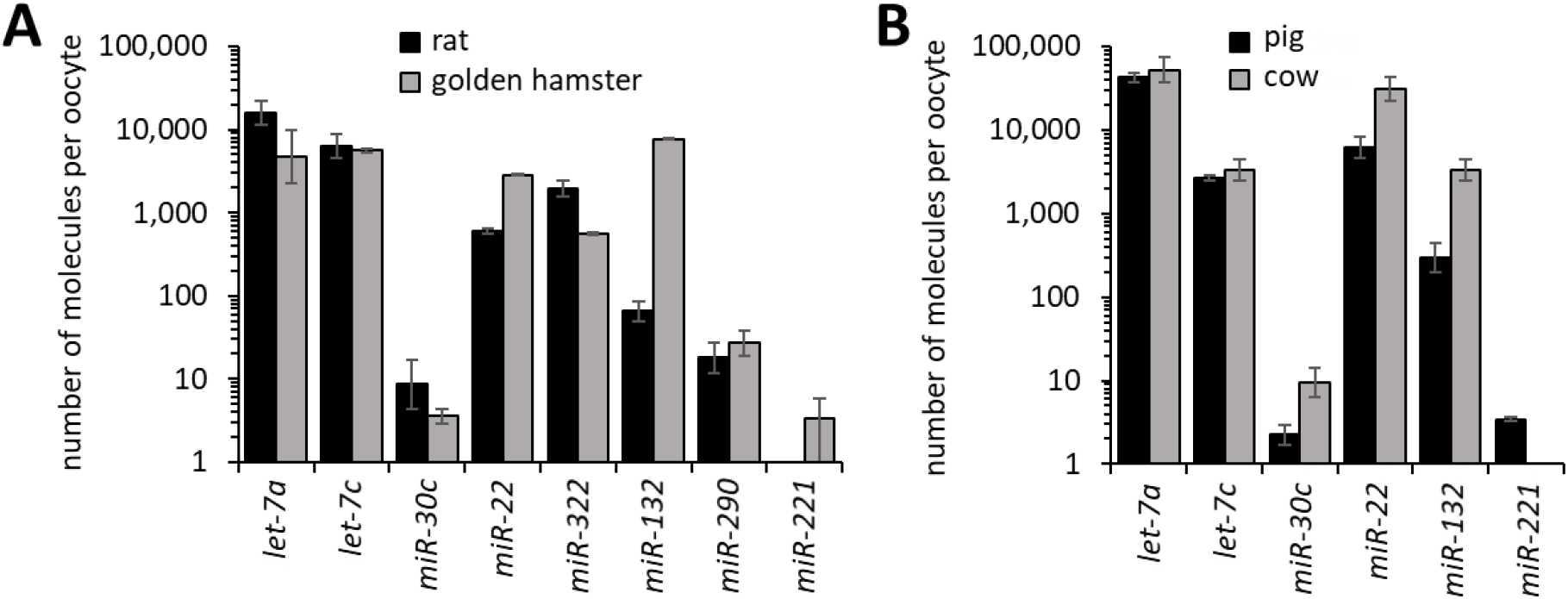
miRNA abundance in various mammalian oocytes. (A) qPCR-based miRNA quantification in rat oocytes (Ø 80 μm) and golden hamster oocytes (Ø 80μm). (B) PCR-based miRNA quantification in porcine oocytes (Ø 105 μm) and bovine oocytes (Ø 120 μm). miRNA copy numbers per cell were estimated in three independent experiments using the standard curve shown in Fig. S1A. Error bars = SD.

### Canonical miRNA-mediated repression is negligible in mammalian oocytes

Previous analysis of miRNA activity in mouse oocytes used firefly and *Renilla* luciferase reporter assays^22^. These reporters were designed^13, 42^ to distinguish the two distinct modes of miRNA-mediated repression (Fig. 3A): (i) RNAi-like endonucleolytic cleavage of perfectly complementary binding sites by AGO2, which was shown previously by mapping cleavage sites^17^, and (ii) translational repression mediated by partial complementarity between a small RNA and its target (typical for animal miRNAs).

**Figure 3.**
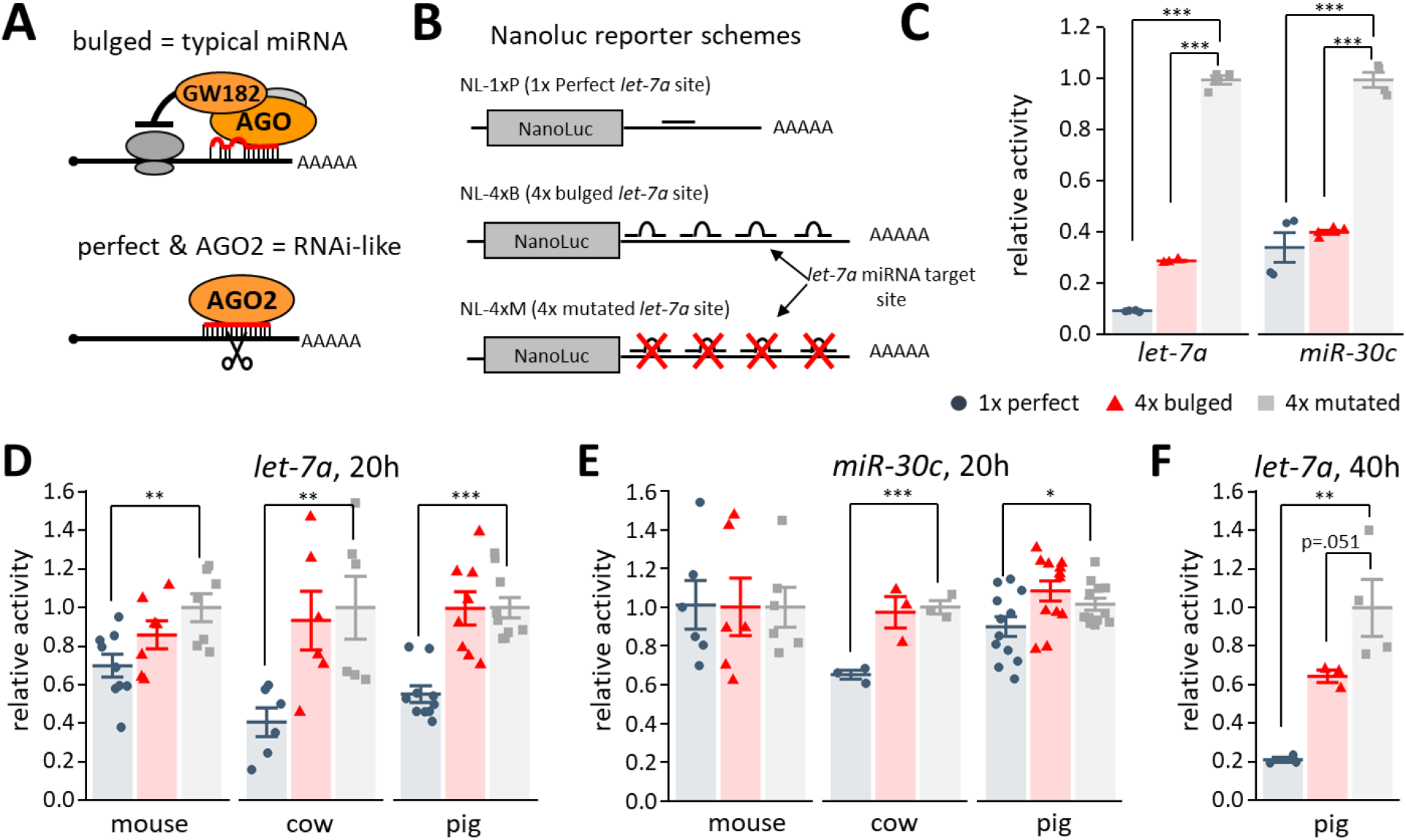
Detection of miRNA activity in mammalian oocytes. (A) Schematic difference between “bulged” and “perfect” miRNA binding sites. Bulged miRNA binding sites, which are typical for animal miRNAs, have imperfect complementarity, and lead to translational repression followed by deadenylation and mRNA degradation^1^. Perfect complementarity of miRNAs loaded on AGO2 results in RNAi-like endonucleolytic cleavage. (B) Schematic depiction of *let-7a* nanoluciferase reporter constructs used in the study. The miRNA sites were cloned in the 3’UTR-either as a 1x-perfect match site, a 4x site in bulged conformation or 4x site mutated. The reporters were *in vitro* transcribed, capped and polyadenylated. A similar set was produced for *miR-30c*. (C) *Let-7a* and *miR-30c* NanoLuc luciferase reporter activities in 3T3 cells. Data represent an average of three independent transfections. (D) Luciferase assay in fully-grown GV mouse, porcine, and bovine oocytes. 10,000 molecules of NanoLuc luciferase reporters carrying 1x-perfect, 4x-bulged or 4x-mutant *let-7a* binding sites were microinjected into GV oocytes, reporter expression was assayed after 20 hours of culture in a medium preventing resumption of meiosis. Error bars = SD. (E) Endogenous miRNA activity repressing bulged *let-7* reporter can be observed in porcine oocytes 40 hours after microinjection (F) *miR-30c* NanoLuc luciferase reporter activity assay in mouse, porcine, and bovine oocytes in three independent experiments. All luciferase data are a ratio of the NanoLuc luciferase reporter activity over a co-injected control firefly luciferase activity. All error bars represent SD. All luciferase data are presented as a ratio of the NanoLuc luciferase signal scaled to coinjected non-targeted firefly luciferase activity, 4x-mutant reporter was set to one. Asterisks indicate statistical significance (p-value) of one-tailed t-test, (* < 0.05, ** < 0.01, *** < 0.001).

The luciferase reporter approach used for studying miRNA activity in oocytes required ~100,000 reporter molecules injected per oocyte^22^, which is in the range of the few most abundant maternal mRNAs^43^. At this level, reporters could have low sensitivity for miRNA-mediated repression. For example, 5,000 repressed reporter molecules would represent 5% of the injected reporter, which would be hardly detectable in this experimental setup. To address this issue, we repeated analysis of miRNA activity with NanoLuc luciferase^44^, a new type of luciferase reporter system, which allowed to reduce the number of reporter molecules by an order of magnitude (10,000 molecules per oocyte) and reliably detect suppression of several thousand reporter mRNA molecules. Furthermore, this amount is within the range of medium-abundant mRNAs such as *Plat* or *Mos*, which were efficiently knocked down by 10,000 long double-stranded RNA molecules per oocyte^45^.

Accordingly, we produced and validated NanoLuc luciferase reporters carrying either a perfectly complementary or four bulged sites to *let-7a* or *miR-30c* miRNAs (Fig. 3B, C). It should be noted that a relative targeted reporter activity is not a precise measure of activity of a specific miRNA used to design binding sites in the targeted reporter. An observed relative reporter repression reflects activity of all miRNAs that suppress the targeted reporter but not its mutated control version. This may involve, for example, other miRNA family members with highly similar sequences, particularly the seed, which may represent the minimum requirement for binding to a cognate RNA^10, 11^.

When we examined reporter repression in oocytes upon injection of 10,000 molecules of NanoLuc reporters, *let-7a* perfect reporter was significantly repressed in mouse oocytes as expected for dominating multiple turnover RNAi-like cleavage (Fig. 3D). Low, albeit statistically insignificant, repression was observed for *Let-7a* bulged reporter (Fig. 3D). *Let-7a* mediated repression was also examined in porcine and bovine oocytes; the bulged reporters did not show any repression while the reporter carrying a binding site perfectly complementary to *let-7a* was repressed by 45% and 60%, respectively (Fig. 3D). On the other hand, the *miR-30c* perfect reporter was not repressed in mouse oocytes (Fig. 3E). This differed from the earlier study^22^ but agreed with low *miR-30c* abundance estimated by qPCR data (Fig. 1A). In cow and pig oocytes, which also have low levels of *miR30c*, we observed low but detectable repression of the *miR-30c* perfect reporter.

Notably, when the porcine oocytes were by chance cultured for 40 hours, which exceeds the time window when GV oocytes maintain their developmental competence^46^, the bulged reporter was repressed by 36% although the difference is not statistically significant (p-value = 0.0506, Fig. 3F). Taken together, reporter results suggested miRNAs in mammalian oocytes do repress their targets; their activity can be detected with perfect reporters but is not high enough to efficiently repress targets with typical miRNA binding sites.

### Increased miRNA levels are sufficient to restore miRNA-mediated repression

Since unfavorable miRNA stoichiometry was observed in all examined mammalian oocytes, we tested if increased miRNA abundance could restore repressive potential of the pathway. This was directly tested by injecting *let-7a* pre-miRNA molecules into mouse (Fig. 4A) and porcine oocytes (Fig. 4B). We observed that 500,000 and 250,000 pre-miRNA molecules were sufficient to restore miRNA pathway activity in oocytes of both species as monitored by repression of bulged reporters. In mouse oocytes, which are smaller than porcine oocytes, we observed repression of bulged reporters even with 100,000 pre-miRNA molecules. This suggests that miRNA abundance in mouse oocytes is not that far from the threshold for significant regulation of the maternal transcriptome.

**Figure 4.**
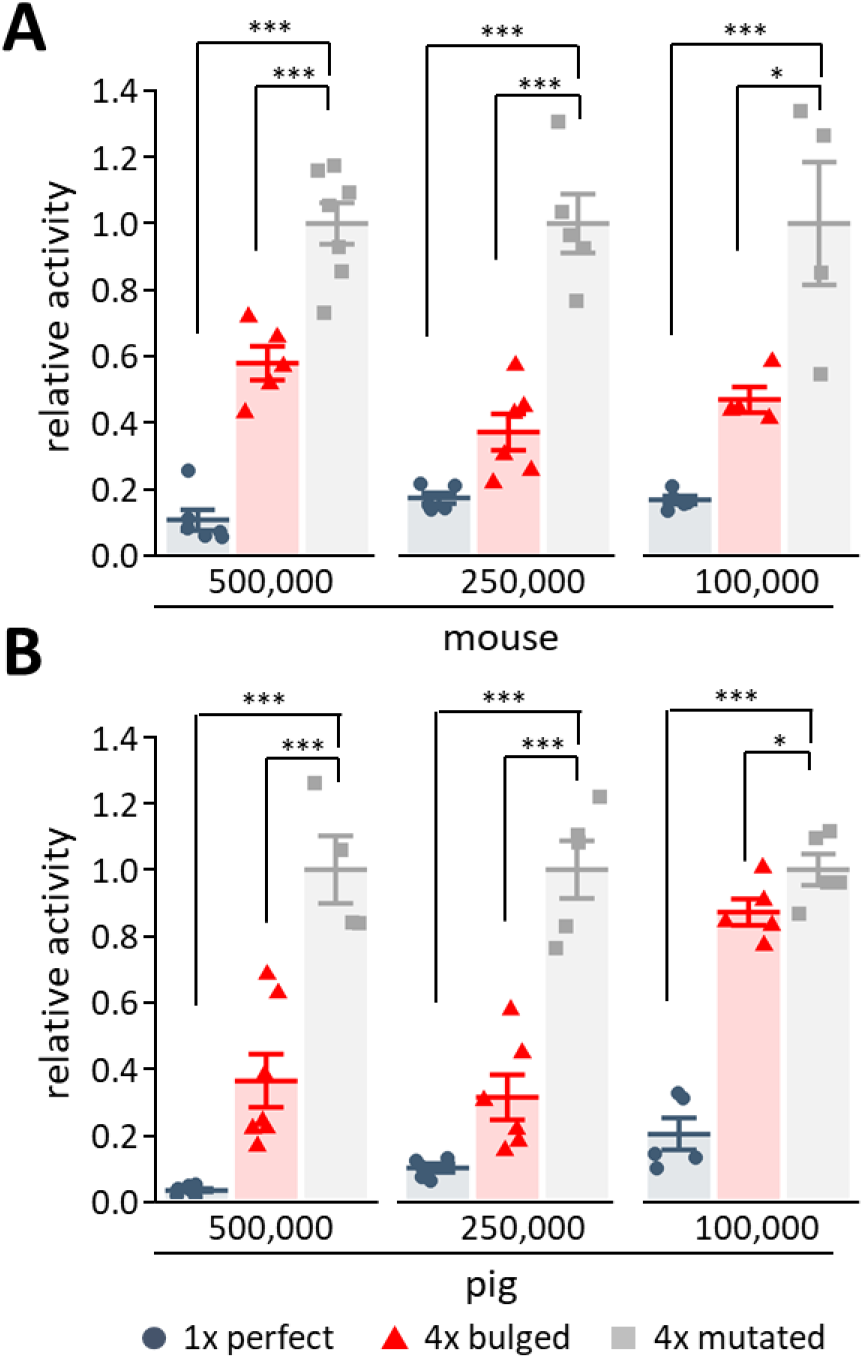
Restoration of miRNA pathway activity in mammalian oocytes. Luciferase reporter activities upon coinjection of mouse (A) or porcine (B) fully-grown GV oocytes of 5×10^5^, 2.5×10^5^ and 1×10^5^ molecules of *let-7a* mimic. The experiment was performed four times. Luciferase data are a presented as a ratio of the NanoLuc luciferase signal scaled to coinjected non-targeted firefly luciferase activity, 4x-mutant reporter was set to one. Error bars = SD. Asterisks indicate statistical significance (p-value) of one-tailed t-test, (* < 0.05, ** < 0.01, *** < 0.001).

Taken together, restored repression of the bulged reporter implies that skewed miRNA:mRNA stoichiometry in oocytes is to a large extent responsible for the apparent lack of miRNA pathway activity. In other words, the miRNA pathway activity is not lost in oocytes, it appears to be present all the time – but the reduced miRNA:mRNA stoichiometry minimizes the significance of miRNA repression.

### Mathematical modeling supports significance of observed miRNA levels

To further examine plausibility of the stoichiometric explanation, we performed computational simulations of target repression using kinetic parameters of target binding and repression derived from experimental analysis of miRNAs^10, 11^ (Fig. 5A). The model simulated miRNA-mediated repression at different concentrations of targets and miRNAs. We assumed no transcription, which is actually the case in fully-grown GV oocytes^47^. The binding of a miRNA to a target was modelled as a one step process using K_D_ 26 pM, which applies for seed-only miRNA:mRNA interaction^11^. Typical miRNA-mediated repression is a multistep process with unknown rate constant(s). Therefore, we replaced typical miRNA-mediated repression with RNAi-like cleavage for which the k_cat_ was measured^11^. The model was then adjusted by a variable coefficient representing strength of miRNA repression relative to RNAi-like cleavage.

**Figure 5.**
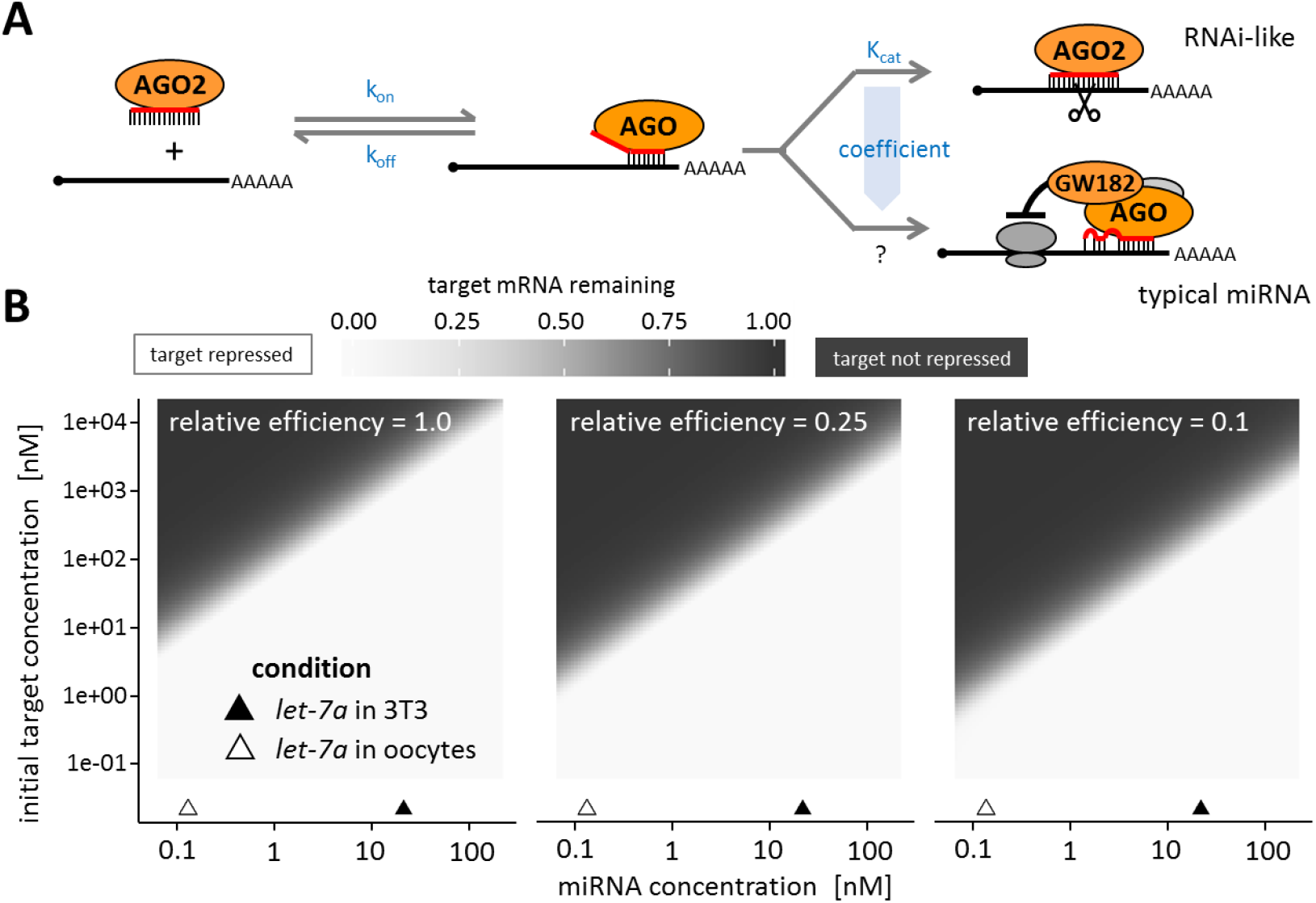
Mathematical simulation of mRNA repression in mouse oocytes. (A) A scheme of the miRNA pathway and parameters used for the mathematical simulation. (B) Mathematical simulation showing proportion of mRNA target repression after 20 hrs. Three different simulation results are shown for different values of efficiency of typical miRNA-mediated repression relative to RNAi-like cleavage. Left: assuming miRNA repression is as effective as RNAi-like cleavage (relative efficiency = 1). Center and right: assuming four and ten times less effective typical miRNA-mediated repression than RNAi-like cleavage (relative efficiency = 0.25 and 0.1, respectively).

For a large set of parameters, the model predicted fast mRNA degradation when the miRNA concentration was similar to the measured concentration of *let-7a* in 3T3 cells while minimal if any degradation was predicted for miRNA concentrations of *let-7a* in oocytes (Fig. 5B, see also Table 1 for concentration calculations). The model predicted that subnanomolar miRNA concentrations observed in fully-grown oocytes could be inefficient for repression of targets above one nM. The entire maternal transcriptome is ~150 nM; if a miRNA would target 5% of the transcriptome through a seed interaction (*let-7* has 1074 predicted mRNAs with 15,028 binding sites^48^), it’s target pool concentration could be just below 10 nM. Taken together, the mathematical modelling demonstrates the significance of two-orders of magnitude difference in concentration of miRNAs in somatic cells and fully-grown oocytes despite the mathematical modelling had to introduce a variable coefficient to address the fact that the miRNA-mediated repression is a multistep process with unknown rate constant(s).

## DISCUSSION

Previous studies showed inefficient miRNA-mediated repression of reporters and no transcriptome changes in mutant oocytes lacking a miRNA biogenesis factor *Dgcr8*, the key component of miRNA biogenesis^22, 23^. Our data show that the loss of miRNA-mediated repression can be largely attributed to the lack of accumulation of maternal miRNAs during oocyte growth resulting in miRNA:mRNA stoichiometry, which impedes efficient miRNA-mediated repression in the fully-grown oocyte. Because some reporter repression could be detected after longer periods (Fig. 3F) or upon increased abundance of a targeting miRNA (Fig. 4), it is possible that other factors, such as reduced deadenylation and decapping^24, 49, 50^, which participate in degrading miRNA-targeted mRNAs, could contribute to the absence of transcriptome changes in *Dgcr8^—/—^* oocytes.

The reason why the number of maternal miRNA molecules does not proportionally increase during the mouse oocyte growth like maternal mRNAs remains unknown. miRNA biogenesis is apparently intact as evidenced by detection of mature miRNAs and undetectable levels of miRNA precursor intermediates. Maternal mRNA acquire extremely extended half-life^31, 32, 33^, which facilitates their accumulation. Post-transcriptional miRNA turnover is regulated in a number of ways but separately from mRNAs (reviewed in^51^). Therefore, the simplest explanation of our data is that miRNA turnover remains unaffected during oogenesis. We propose that miRNAs did not evolve a specific mechanism facilitating their accumulation. Consequently, their steady state amount would not change during oocyte growth but cytoplasmic concentration would dive two orders of magnitude as the oocyte would grow into a diameter of 80 μm.

It was also proposed that the non-functionality of miRNAs in mouse oocytes is associated with an oocyte-specific truncated *Ago2* variant, which reduces the expression of the full-length functional *Ago2*^25^. Limiting amount of AGO2 could explain reduced miRNA levels. However, this alternative processing of *Ago2* transcript is not observed in other examined mammalian oocytes whose transcriptomes were sequenced^40, 41^. Thus, other mammals would have to independently evolve other mechanism(s) restricting the miRNA pathway via expression of AGO proteins. Alternatively, mouse AGO2 regulation reported by Freimer et al.^25^ could be an adaptation restricting the endogenous RNAi pathway, which is highly active in mouse oocytes and has been specifically evolving in the mouse lineage^37, 41, 52^.

We previously reported disappearance of processing bodies (P-bodies) during oocyte growth^24^. P-bodies are cytoplasmic foci forming as liquid–liquid phase separation from proteins involved in RNA metabolism, miRNA effector complexes and their targets (reviewed in^53^). We initially hypothesized that P-body disappearance may reflect inhibition of the miRNA pathway, possibly caused by some defect in formation of the effector complex. However, in the light of the current data, disappearance of P-bodies could be plausibly explained by dilution caused by oocyte growth.

Importantly, we show that increased level of miRNA precursor is sufficient to induce significant repression of bulged NanoLuc reporters. This suggest that miRNA processing has sufficient capacity and more miRNAs could be accommodated on AGO2 in porcine and mouse oocytes. Of note is that Freimer et al. did not observe repression of the bulged reporter upon injection of miRNA precursors into mouse oocytes^25^. However, the *Renilla*/Firefly luciferase reporter system requires an order of magnitude more reporter molecules than our NanoLuc luciferase system and hence may be insensitive to repression of several thousand reporter mRNAs. Therefore, we think that our reporter results do not contradict those of Freimer et al. but rather complement them as we developed a quantitative framework for explaining the lack of functional significance of maternal miRNAs.

It is surprising that such an omnipresent post-transcriptional regulatory pathway would lose its repressive potential in mammalian oocytes and play no significant role in the mammalian transition from maternal to zygotic transcriptome control. There are several hypothetical scenarios, which could be examined in future studies. First, there may be minimal pressure to stabilize maternal gene expression program in non-dividing oocytes through miRNAs. Second, retaining a functional miRNA pathway may not be compatible or may even be detrimental for mechanisms, which oocytes employ for maternal-to-zygotic (MZT) transition. This could concern post-transcriptional control of the maternal transcriptome such as storage of deadenylated dormant maternal mRNAs or accumulation of totipotency factors in the presence of *let-7* miRNA (reviewed in^54^). Third, the lack of miRNA accumulation could be adopted as a strategy for replacing maternal miRNAs with pluripotent zygotic miRNAs of the *miR-290* family, where *let-7* could be seen as an anti-pluripotency factor^55^. This notion is consistent with miRNA activity during MZT in zebrafish where zygotic miRNAs have dominant functional significance in the transition^56^.

Taken together, we show that the miRNA pathway is disabled in different mammalian oocytes. The inability of maternal miRNAs to exert significant mRNA repression stems from skewed stoichiometry. Increased miRNA concentration is sufficient to restore target repression. This demonstrates that the miRNA pathway activity in oocytes is maintained not too far from a threshold for effective repression. This probably facilitates bringing the miRNA pathway back into biologically relevant role upon zygotic genome activation in the cleaving preimplantation embryo.

The relatively passive mechanism by which mammalian oocytes would abandon the miRNA pathway implies lack of positive selection for miRNA function in growing and fully-grown oocytes. Consequently, miRNA metabolism would not adapt to the accumulation of maternal mRNAs during oocyte’s growth. While we show that this is a common mammalian phenomenon, the lack of miRNA activity was also observed in frog oocytes^25^ and miRNA depletion seems to exist in zebrafish oocyte as well^57^. Therefore, this phenomenon appears to be a common feature of vertebrate oocytes and we speculate that it could exist in many other taxons.

## MATERIAL AND METHODS

### Oocyte collection and microinjection

Animal experiments were approved by the Institutional Animal Use and Care Committees (approval no. 34/2014) and were carried out in accordance with the European Union regulations. C57BI/6NCrl mice were obtained from Charles Rivers. Meiotically incompetent oocytes were collected from 8-12 days old females whereas fully-grown GV oocytes were isolated from 7-9 weeks old females. Golden hamsters were obtained from Charles Rivers. Oocytes were collected from ovaries of freshly sacrificed 12 month old animals. Porcine and bovine oocytes were obtained from slaughterhouse material as described previously^58, 59^.

Meiotically incompetent oocytes were isolated by incubating the ovaries in 1XPBS with 1 mg/ml collagenase (Sigma) at 37 °C and then were collected in M2 medium (Sigma). Fully-grown GV oocytes and hamster oocytes were collected by puncturing the antral follicles with syringe needles followed by collection in M2 medium containing 0.2 mM isobutylmethylxanthine (IBMX; Sigma) to prevent resumption of meiosis.

For microinjection, a mixture of *in-vitro* transcribed firefly and nanoluciferase RNA in the ratio of 100,000:10,000 molecules with or without *let-7a* mimic (Sigma) was injected per oocyte with an Eppendorf FemtoJet. Mouse oocytes post injection were cultured in M16 media (Merck) supplemented with IBMX in 5% CO_2_ at 37 °C for 20 hours. Bovine oocytes were cultured in MPM media (prepared in house) supplemented with milrinone (Sigma) without a paraffin overlay in a humidified atmosphere at 39 °C with 5% CO_2_ for 20 h. Pig oocytes were cultured in M-199 MEDIUM (Gibco) supplemented with 1 mM dbcAMP, 0.91 mM sodium pyruvate, 0.57 mM cysteine, 5.5 mM Hepes, antibiotics and 5% fetal calf serum (Sigma). Injected oocytes were incubated at 38.5 °C in a humidified atmosphere of 5% CO_2_ for 20 h.

### cDNA synthesis and qPCR

The oocytes were washed with M2 media to remove any residual IBMX and collected in minimum amount of M2 media with 1 ul of Ribolock and incubated at 85 °C for 5 minutes to release the RNA. NIH 3T3 cells were counted and then lysed using Real time ready cell lysis kit (Roche). cDNA synthesis for miRNA was done with the miRCURY LNA RT kit (Qiagen) according to the manufacturer’s protocol. qPCR reaction was set using single oocyte per well with the miRCURY LNA SYBR green kit (Qiagen) as per manufacturers protocol in a Roche LightCycler 480. miRCURY LNA miRNA specific primers were used and are listed in the resource table. The standard curve reactions were carried out using *mmu-miR-221* and *let-7a* oligonucleotides obtained from Integrated DNA technologies (see Table S2 for oligonucleotide sequences).

A standard curve was first plotted using *let-7a* mimic. The oligonucleotide was either first serially diluted and then cDNA prepared for each dilution (Fig. S1A) or the cDNA was prepared first and then serially diluted (Fig. S1B) to rule out the variability in the efficiency of cDNA synthesis. Standard curve was also standardized using *miR-221* mimic as it was not detected in fully-grown mouse oocyte (Fig. 1A). It therefore could be added to fully-grown mouse oocyte lysate and then used to set cDNA to avoid any discrepancies that might occur as a consequence of additional oocyte specific factors (Fig. S1C).

### Northern blot

The sequence for *let-7a* standard used for northern blotting was the same as mentioned above but with 5’P. Oocytes were collected in 1XPBS with Ribolock and the RNA was isolated with Trizol. Northern blot probes were labelled by incubating the RNA with 20 μCi of γ-^32^P-ATP (Hartmann Analytic) and 0.5 U/μl T4 Polynucleotide Kinase (PNK) in 1x PNK buffer A (Thermo Scientific) at 37 °C for 30-60 min. The reaction was stopped by adding EDTA, pH 8.0, and the labelled oligonucleotides were purified with a G-25 column (GE Healthcare). RNAs were separated in 1x TBE on a 12% polyacrylamide urea gel. The RNA was then blotted for 30 min at 20 V onto an Amersham Hybond-N membrane (GE Healthcare) and crosslinked to the membrane for 1 h at 50°C using a 1-ethyl-3-(3-dimethylaminopropyl) carbodiimide-solution. The membrane was incubated over night at 50 °C in hybridization solution (5x SSC, 7% SDS, 20 mM sodium phosphate buffer pH 7.2, 1x Denhardt’s solution) with a ^32^P-labelled oligonucleotide antisense to the small RNA. The membrane was washed twice with 5x SSC, 1% SDS, once with 1x SSC, 1% SDS and wrapped in saran. Signals were detected by exposure to a storage phosphor screen and scanning with the Personal Molecular Imager (Bio-Rad). For further details refer to^60^

### Plasmid reporters

Nanoluciferase plasmid pNL1.1 was obtained from Promega. Nluc gene was cleaved out from this plasmid and ligated in the phRL-SV40 plasmid downstream of the T7 promoter. miRNA binding sites were ordered as oligos from Sigma. Oligos were annealed, phosphorylated and then cloned downstream of Nluc gene in XbaI site. The firefly control plasmid was made in house with a T7 promoter (see Table S2 for oligonucleotide sequences-restriction overhangs underlined).

### Cell lines and cell culture

NIH 3T3 mouse fibroblast cells were grown in DMEM supplemented with 10% Fetal Bovine Serum and transfected with miRNA-targeted reporters as described previously^22^.

### *In vitro* transcription

Linearized pNL-Let7a-perfect, bulged, mutant and Firefly plasmids were *in vitro* transcribed, capped and then polyadenylated by Poly(A) tailing kit (ThermoFisher).

### Luciferase assay

5 oocytes were collected per aliquot and Nano-Glo Dual-Luciferase reporter assay system was used to measure the samples as per the manufacturer’s protocol (Promega).

### Mathematical modelling

The mathematical model for miRNA degradation introduces the following simplifying assumptions:

- miRNA degradation is modelled as a one step process (similar to RNAi pathway), but with lower catalytic efficiency than the RNAi pathway.
- The AGO2 concentration is assumed to not be rate limiting, i.e. all miRNAs are assumed to be loaded on AGO2 at any tested concentration.
- The concentration of AGO2 + miRNA complex is assumed to be constant, i.e. there is neither synthesis or degradation of miRNA.
- There is no target mRNA synthesis.
- All assumed miRNA binding events are to be those of pure seed pairing.
- Only uncleaved free mRNA is considered in simulations (mRNA bound in complex with AGO2 + miRNA is not considered).

Those assumptions let us describe the system as a set of differential equations, identical to the ones used in Michaelis-Menten kinetics:

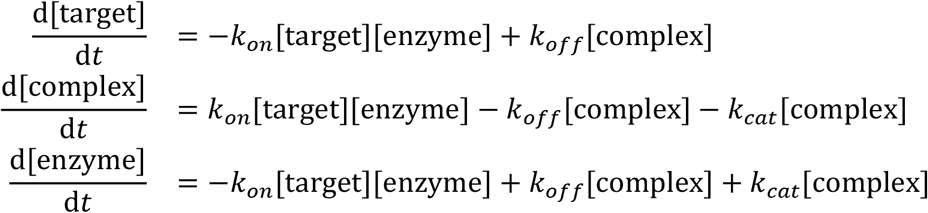

Where *k*_*on*_,*k*_*off*_ and *k*_*cat*_ are the reaction rates in the system and [target], [complex] and [enzyme] are the concentrations of free mRNA molecules with target sites for the miRNA in question, mRNA + miRNA + AGO2 complex and miRNA + AGO2 complex respectively. The simulation was written in the R language (https://www.R-project.org/) using packages deSolve^61^ and visualised via ggplot2^62^. More details about the simulation, including full source code is given in the supplementary material.

## ACKNOWLEDGEMENTS

We would like to thank Anna Tetkova and Ales Petelak for help with establishing microinjection experiments and Tatiana Spitzova and Jovana Sadlova for assistance with hamster oocyte isolation. This work was funded from the European Research Council under the European Union’s Horizon 2020 research and innovation programme (grant agreement No 647403, D-FENS). Additional support was provided by the Ministry of Education, Youth, and Sports (MEYS) project NPU1 LO1419. Work in the Meister lab was supported by the Deutsche Forschungsgemeinschaft (SFB960). J.K. and V.K. were in part supported by IAPG institutional support RVO: 67985904. M.M. was supported by the ELIXIR_CZ research infrastructure MEYS project no. LM2015047. Financial support of S.K. was in part provided by the Charles University in a form of a PhD student fellowship; this work will be in part used to fulfil requirements for a PhD degree and hence can be considered “school work”.

## AUTHOR CONTRIBUTIONS

S.K., V.K. and D.Z. performed the experiments. M.M. did the mathematical modelling. S.K, R.M., J.K., G.M. and P.S. designed the experiments, supervised, and wrote the manuscript.

## SUPPLEMENTAL INFORMATION

**Figure S1.**
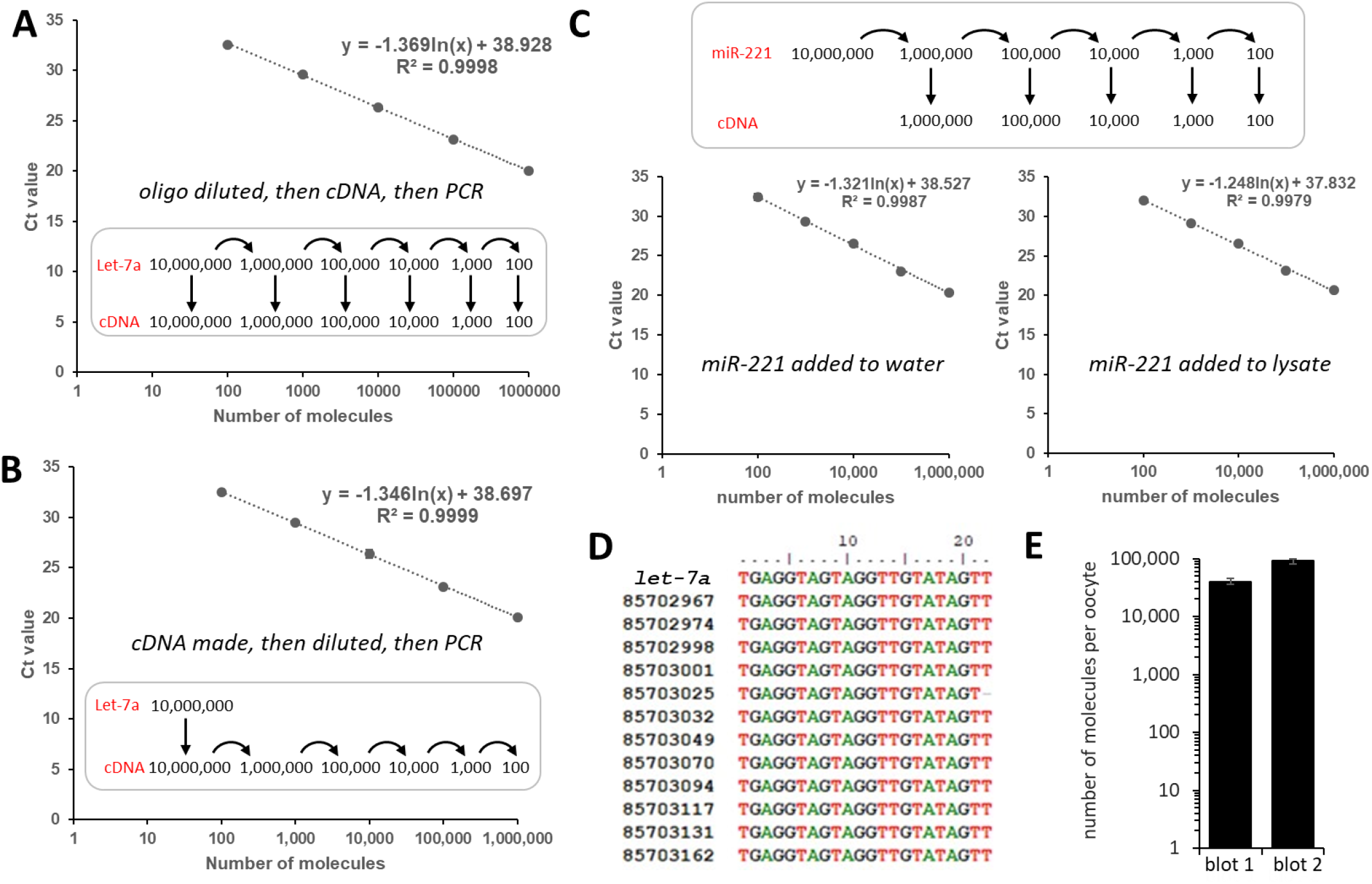
Supplementary data for miRNA quantification. (A) Calibration curve made from a *let-7a* synthetic single-stranded RNA oligonucleotide. The oligonucleotide was serially diluted first and then cDNA was set for each dilution followed by qPCR. (B) Calibration curve made from a *let-7a* oligonucleotide where the oligonucleotide was first reverse transcribed into cDNA, then the cDNA was serially diluted for qPCR. (C) Calibration curve made from a *miR-221* oligo. The oligonucleotide was serially diluted first and then reverse transcription was done or each its dilution was added to a mouse oocyte lysate before reverse transcription (*miR-221* was undetectable in mouse oocytes). A-C calibrations were performed three times, data points represent average values. (D) Alignment of sequences of twelve subcloned *let-7a* qPCR products. (E) Quantification of *let-7a* northern blots (Fig. 1B).

**Supplemental Table S1.**
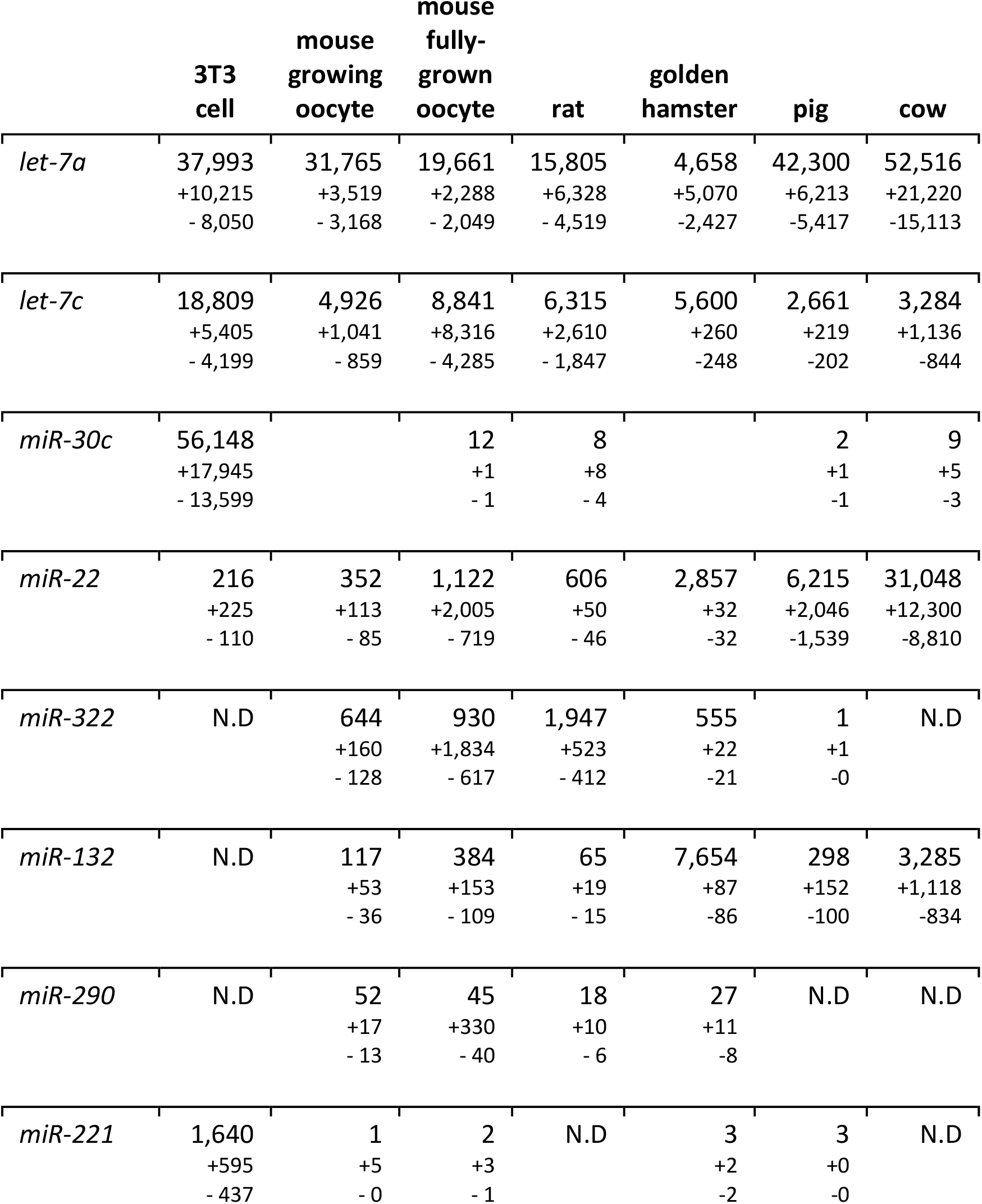
Quantifications of miRNA molecules per cell (+/- SD)

|  | **3T3**<br>**cell** | **mouse**<br>**growing**<br>**oocyte** | **mouse**<br>**fully-**<br>**grown**<br>**oocyte** | **rat** | **golden**<br>**hamster** | **pig** | **cow** |
| --- | --- | --- | --- | --- | --- | --- | --- |
| *let-7a* | 37,993<br>+10,215<br>- 8,050 | 31,765<br>+3,519<br>- 3,168 | 19,661<br>+2,288<br>- 2,049 | 15,805<br>+6,328<br>- 4,519 | 4,658<br>+5,070<br>-2,427 | 42,300<br>+6,213<br>-5,417 | 52,516<br>+21,220<br>-15,113 |
| *let-7c* | 18,809<br>+5,405<br>- 4,199 | 4,926<br>+1,041<br>- 859 | 8,841<br>+8,316<br>- 4,285 | 6,315<br>+2,610<br>- 1,847 | 5,600<br>+260<br>-248 | 2,661<br>+219<br>-202 | 3,284<br>+1,136<br>-844 |
| *miR-30c* | 56,148<br>+17,945<br>- 13,599 |  | 12<br>+1<br>- 1 | 8<br>+8<br>- 4 |  | 2<br>+1<br>-1 | 9<br>+5<br>-3 |
| *miR-22* | 216<br>+225<br>- 110 | 352<br>+113<br>- 85 | 1,122<br>+2,005<br>- 719 | 606<br>+50<br>- 46 | 2,857<br>+32<br>-32 | 6,215<br>+2,046<br>-1,539 | 31,048<br>+12,300<br>-8,810 |
| *miR-322* | N.D | 644<br>+160<br>- 128 | 930<br>+1,834<br>- 617 | 1,947<br>+523<br>- 412 | 555<br>+22<br>-21 | 1<br>+1<br>-0 | N.D |
| *miR-132* | N.D | 117<br>+53<br>- 36 | 384<br>+153<br>- 109 | 65<br>+19<br>- 15 | 7,654<br>+87<br>-86 | 298<br>+152<br>-100 | 3,285<br>+1,118<br>-834 |
| *miR-290* | N.D | 52<br>+17<br>- 13 | 45<br>+330<br>- 40 | 18<br>+10<br>- 6 | 27<br>+11<br>-8 | N.D | N.D |
| *miR-221* | 1,640<br>+595<br>- 437 | 1<br>+5<br>- 0 | 2<br>+3<br>- 1 | N.D | 3<br>+2<br>-2 | 3<br>+0<br>-0 | N.D |

**Supplemental Table S2.**
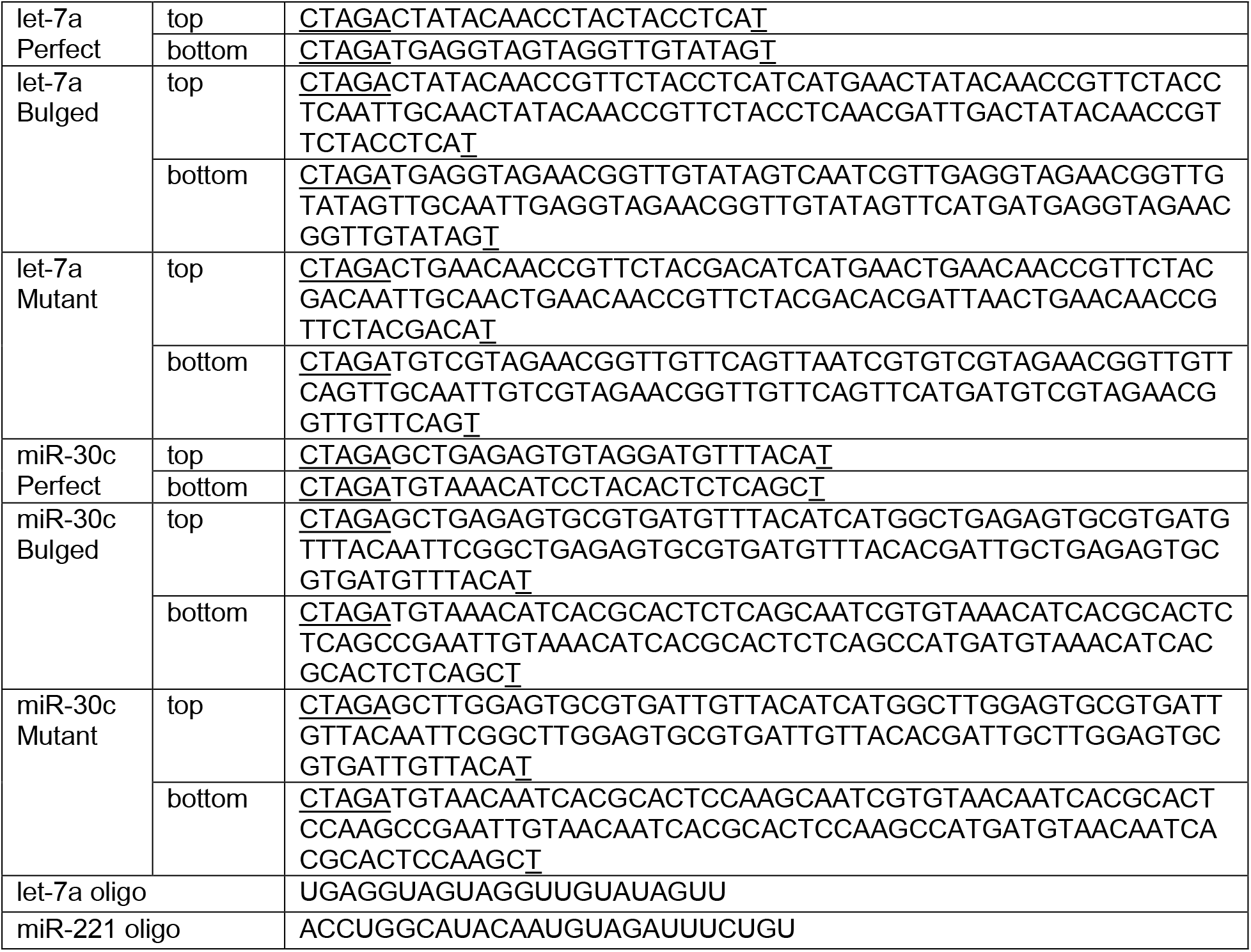
Oligonucleotide table.

### Supplementary File

**2019 Kataruka et al. supplementary_code.zip**

The code used for mathematical modeling

## REFERENCES

1. Bartel DP. Metazoan MicroRNAs. Cell 173, 20–51 (2018).

2. Dueck A, Meister G. Assembly and function of small RNA - argonaute protein complexes. Biol Chem 395, 611–629 (2014).

3. Bosson AD, Zamudio JR, Sharp PA. Endogenous miRNA and target concentrations determine susceptibility to potential ceRNA competition. Mol Cell 56, 347–359 (2014).

4. Denzler R, Agarwal V, Stefano J, Bartel DP, Stoffel M. Assessing the ceRNA hypothesis with quantitative measurements of miRNA and target abundance. Mol Cell 54, 766–776 (2014).

5. Denzler R, McGeary SE, Title AC, Agarwal V, Bartel DP, Stoffel M. Impact of MicroRNA Levels, Target-Site Complementarity, and Cooperativity on Competing Endogenous RNA-Regulated Gene Expression. Mol Cell 64, 565–579 (2016).

6. Bishop JO, Morton JG, Rosbash M, Richardson M. Three abundance classes in HeLa cell messenger RNA. Nature 250, 199–204 (1974).

7. Yekta S, Shih IH, Bartel DP. MicroRNA-directed cleavage of HOXB8 mRNA. Science 304, 594–596 (2004).

8. Brennecke J, Stark A, Russell RB, Cohen SM. Principles of microRNA-target recognition. PLoS Biol 3, e85 (2005).

9. Sontheimer EJ. Assembly and function of RNA silencing complexes. Nat Rev Mol Cell Biol 6, 127–138 (2005).

10. Salomon WE, Jolly SM, Moore MJ, Zamore PD, Serebrov V. Single-Molecule Imaging Reveals that Argonaute Reshapes the Binding Properties of Its Nucleic Acid Guides. Cell 162, 84–95 (2015).

11. Wee LM, Flores-Jasso CF, Salomon WE, Zamore PD. Argonaute divides its RNA guide into domains with distinct functions and RNA-binding properties. Cell 151, 1055–1067 (2012).

12. Doench JG, Petersen CP, Sharp PA. siRNAs can function as miRNAs. Genes Dev 17, 438–442 (2003).

13. Hutvagner G, Zamore PD. A microRNA in a multiple-turnover RNAi enzyme complex. Science 297, 2056–2060 (2002).

14. Lim LP, et al. Microarray analysis shows that some microRNAs downregulate large numbers of target mRNAs. Nature 433, 769–773 (2005).

15. Kozomara A, Birgaoanu M, Griffiths-Jones S. miRBase: from microRNA sequences to function. Nucleic Acids Res 47, D155–D162 (2019).

16. Krutzfeldt J, et al. Silencing of microRNAs in vivo with ‘antagomirs’. Nature 438, 685–689 (2005).

17. Schmitter D, et al. Effects of Dicer and Argonaute down-regulation on mRNA levels in human HEK293 cells. Nucleic Acids Res 34, 4801–4815 (2006).

18. Wang Y, Medvid R, Melton C, Jaenisch R, Blelloch R. DGCR8 is essential for microRNA biogenesis and silencing of embryonic stem cell self-renewal. Nature genetics 39, 380–385 (2007).

19. Friedman RC, Farh KK, Burge CB, Bartel DP. Most mammalian mRNAs are conserved targets of microRNAs. Genome research 19, 92–105 (2009).

20. Tam OH, et al. Pseudogene-derived small interfering RNAs regulate gene expression in mouse oocytes. Nature 453, 534–538 (2008).

21. Watanabe T, et al. Endogenous siRNAs from naturally formed dsRNAs regulate transcripts in mouse oocytes. Nature 453, 539–543 (2008).

22. Ma J, et al. MicroRNA activity is suppressed in mouse oocytes. Curr Biol 20, 265–270 (2010).

23. Suh N, et al. MicroRNA function is globally suppressed in mouse oocytes and early embryos. Curr Biol 20, 271–277 (2010).

24. Flemr M, Ma J, Schultz RM, Svoboda P. P-body loss is concomitant with formation of a messenger RNA storage domain in mouse oocytes. Biol Reprod 82, 1008–1017 (2010).

25. Freimer JW, Krishnakumar R, Cook MS, Blelloch R. Expression of Alternative Ago2 Isoform Associated with Loss of microRNA-Driven Translational Repression in Mouse Oocytes. Curr Biol 28, 296–302 e293 (2018).

26. Hastie ND, Bishop JO. The expression of three abundance classes of messenger RNA in mouse tissues. Cell 9, 761–774 (1976).

27. Piko L, Clegg KB. Quantitative changes in total RNA, total poly(A), and ribosomes in early mouse embryos. Dev Biol 89, 362–378 (1982).

28. Carter MG, et al. Transcript copy number estimation using a mouse whole-genome oligonucleotide microarray. Genome Biol 6, R61 (2005).

29. Marinov GK, et al. From single-cell to cell-pool transcriptomes: stochasticity in gene expression and RNA splicing. Genome research 24, 496–510 (2014).

30. Fan X, et al. Single-cell RNA-seq transcriptome analysis of linear and circular RNAs in mouse preimplantation embryos. Genome Biol 16, 148 (2015).

31. Jahn CL, Baran MM, Bachvarova R. Stability of RNA synthesized by the mouse oocyte during its major growth phase. J Exp Zool 197, 161–171 (1976).

32. Brower PT, Gizang E, Boreen SM, Schultz RM. Biochemical studies of mammalian oogenesis: synthesis and stability of various classes of RNA during growth of the mouse oocyte in vitro. Dev Biol 86, 373–383 (1981).

33. De Leon V, Johnson A, Bachvarova R. Half-lives and relative amounts of stored and polysomal ribosomes and poly(A) + RNA in mouse oocytes. Dev Biol 98, 400–408 (1983).

34. Garcia-Lopez J, Hourcade Jde D, Alonso L, Cardenas DB, del Mazo J. Global characterization and target identification of piRNAs and endo-siRNAs in mouse gametes and zygotes. Biochim Biophys Acta 1839, 463–475 (2014).

35. Yang Q, et al. Highly sensitive sequencing reveals dynamic modifications and activities of small RNAs in mouse oocytes and early embryos. Sci Adv 2, e1501482 (2016).

36. Tang F, et al. Maternal microRNAs are essential for mouse zygotic development. Genes Dev 21, 644–648 (2007).

37. Flemr M, et al. A retrotransposon-driven dicer isoform directs endogenous small interfering RNA production in mouse oocytes. Cell 155, 807–816 (2013).

38. Murchison EP, et al. Critical roles for Dicer in the female germline. Genes Dev 21, 682–693 (2007).

39. Kaneda M, Tang F, O’Carroll D, Lao K, Surani MA. Essential role for Argonaute2 protein in mouse oogenesis. Epigenetics Chromatin 2, 9 (2009).

40. Graf A, Krebs S, Zakhartchenko V, Schwalb B, Blum H, Wolf E. Fine mapping of genome activation in bovine embryos by RNA sequencing. Proc Natl Acad Sci U S A 111, 4139–4144 (2014).

41. Franke V, et al. Long terminal repeats power evolution of genes and gene expression programs in mammalian oocytes and zygotes. Genome research 27, 1384–1394 (2017).

42. Pillai RS, et al. Inhibition of translational initiation by Let-7 MicroRNA in human cells. Science 309, 1573–1576 (2005).

43. Roller RJ, Kinloch RA, Hiraoka BY, Li SS, Wassarman PM. Gene expression during mammalian oogenesis and early embryogenesis: quantification of three messenger RNAs abundant in fully grown mouse oocytes. Development 106, 251–261 (1989).

44. England CG, Ehlerding EB, Cai W. NanoLuc: A Small Luciferase Is Brightening Up the Field of Bioluminescence. Bioconjug Chem 27, 1175–1187 (2016).

45. Svoboda P, Stein P, Hayashi H, Schultz RM. Selective reduction of dormant maternal mRNAs in mouse oocytes by RNA interference. Development 127, 4147–4156 (2000).

46. Grupen CG, Fung M, Armstrong DT. Effects of milrinone and butyrolactone-I on porcine oocyte meiotic progression and developmental competence. Reprod Fertil Dev 18, 309–317 (2006).

47. Svoboda P, Franke V, Schultz RM. Sculpting the Transcriptome During the Oocyte-to-Embryo Transition in Mouse. Curr Top Dev Biol 113, 305–349 (2015).

48. Agarwal V, Bell GW, Nam JW, Bartel DP. Predicting effective microRNA target sites in mammalian mRNAs. Elife 4, (2015).

49. Ma J, Flemr M, Strnad H, Svoboda P, Schultz RM. Maternally recruited DCP1A and DCP2 contribute to messenger RNA degradation during oocyte maturation and genome activation in mouse. Biol Reprod 88, 11 (2013).

50. Ma J, Fukuda Y, Schultz RM. Mobilization of Dormant Cnot7 mRNA Promotes Deadenylation of Maternal Transcripts During Mouse Oocyte Maturation. Biol Reprod 93, 48 (2015).

51. Krol J, Loedige I, Filipowicz W. The widespread regulation of microRNA biogenesis, function and decay. Nat Rev Genet 11, 597–610 (2010).

52. Stein P, et al. Essential Role for endogenous siRNAs during meiosis in mouse oocytes. PLoS Genet 11, e1005013 (2015).

53. Luo Y, Na Z, Slavoff SA. P-Bodies: Composition, Properties, and Functions. Biochemistry 57, 2424–2431 (2018).

54. Svoboda P. Why mouse oocytes and early embryos ignore miRNAs? RNA Biol 7, 559–563 (2010).

55. Svoboda P, Flemr M. The role of miRNAs and endogenous siRNAs in maternal-to-zygotic reprogramming and the establishment of pluripotency. EMBO Rep 11, 590–597 (2010).

56. Giraldez AJ, et al. Zebrafish MiR-430 promotes deadenylation and clearance of maternal mRNAs. Science 312, 75–79 (2006).

57. Chen PY, et al. The developmental miRNA profiles of zebrafish as determined by small RNA cloning. Genes Dev 19, 1288–1293 (2005).

58. Blaha M, Nemcova L, Kepkova KV, Vodicka P, Prochazka R. Gene expression analysis of pig cumulus-oocyte complexes stimulated in vitro with follicle stimulating hormone or epidermal growth factor-like peptides. Reprod Biol Endocrinol 13, 113 (2015).

59. Kinterova V, Kanka J, Petruskova V, Toralova T. Inhibition of Skp1-Cullin-F-box complexes during bovine oocyte maturation and preimplantation development leads to delayed development of embryosdagger. Biol Reprod 100, 896–906 (2019).

60. Hasler D, et al. The Lupus Autoantigen La Prevents Mis-channeling of tRNA Fragments into the Human MicroRNA Pathway. Mol Cell 63, 110–124 (2016).

61. Soetaert K, Petzoldt T, Setzer RW. Solving Differential Equations in R: Package deSolve. 2010 33, 25 (2010).

62. Wickham H. ggplot2: Elegant Graphics for Data Analysis. Springer Publishing Company, Incorporated (2009).

