## Supplementary code file for "Low miRNA abundance disables microRNA pathway in mammalian oocytes": miRNA-sim.html

Simple miRNA computational modelling


Code 

- Show All Code
- Hide All Code

### Simple miRNA computational modelling

###### Martin Modrak

#### 2019-07-02

Abstract

This is a supplement to the paper “Low miRNA abundance disables microRNA pathway in mammalian oocytes”. We describe a simplified mathematical model of the miRNA pathway based on our knowledge of the RNAi pathway and show simulation results supporting the conclusions of the paper.

```
#knitr::opts_chunk$set(echo = FALSE)
#Next line is useful for Word output
#knitr::opts_chunk$set(fig.width = 9)
library(deSolve)
library(ggplot2)
library(dplyr)
library(tidyr)
library(tibble)
library(cowplot) #Optional, can be commented out
library(knitr)
options(dplyr.show_progress = FALSE)

temp_dir <- "local_temp_data"
if(!dir.exists(temp_dir)) {
  dir.create(temp_dir)
}
```

*Source for this file is also available at https://github.com/martinmodrak/miRNA-sim*

### Assumptions

- miRNA degradation is modelled as one step process (similar to RNAi pathway), but with lower catalytic efficiency than the RNAi pathways
- The AGO2 concentration is assumed to not be rate limiting, in particular all miRNA is assumed to be loaded on AGO2 at all times
- The concentration of AGO2 + miRNA complex is assumed to be constant.
- All miRNA binding events are assumed to be those of pure seed pairing (for seed + additionial base pairing the binding is slightly stronger).
- Only uncleaved free mRNA is observed in experiments (mRNA bound in complex with AGO2 + miRNA is not be observed).
- No mRNA synthesis is taking place.

### Mathematical model

The assumptions above lead to a simple description of the system as a set of differential equations, identical to the ones used in Michaelis-Menten kinetics. Denoting the concentration of free mRNA molecules with target sites for the miRNA in question as \([\mathrm{target}]\), the concentration of mRNA + miRNA + AGO2 complex as \([\mathrm{complex}]\) and the concentration of the miRNA + AGO2 complex as \([\mathrm{enzyme}]\) The kinetics are descibed as:

\[
\begin{align}
\frac{\mathrm{d}[\mathrm{target}]}{\mathrm{d}t} &= -k\_{on}[\mathrm{target}][\mathrm{enzyme}] + k\_{off}[\mathrm{complex}] \\
\frac{\mathrm{d}[\mathrm{complex}]}{\mathrm{d}t} &= k\_{on}[\mathrm{target}][\mathrm{enzyme}] - k\_{off}[\mathrm{complex}] - k\_{cat}[\mathrm{complex}] \\
\frac{\mathrm{d}[\mathrm{enzyme}]}{\mathrm{d}t} &= -k\_{on}[\mathrm{target}][\mathrm{enzyme}] + k\_{off}[\mathrm{complex}] + k\_{cat}[\mathrm{complex}] \\
\end{align}
\]

Where \(k\_{on}\),\({k\_{off}}\) and \(k\_{cat}\) are the reaction rates in the system.

### Input data

```
### Define units and conversions
#Using litre as volume unit! (since constants are in moles per litre)
mole_unit <- 6.022e14 #nano mole
mole_unit_from_nano_mole <- 1
mole_unit_from_mole <- 1e9
#Using 1/10th second as time unit (some sims need fine resolution)
time_unit_from_sec <- 1

K_M_RNAi <- 0.1 * mole_unit_from_nano_mole
k_on_RNAi <- 3.6e7 / (mole_unit_from_mole * time_unit_from_sec)

k_off_RNAi <- 7.7e-4 / time_unit_from_sec
k_cat_RNAi <- 8.1e-4 / time_unit_from_sec
k_cat_RNAi_computed <- K_M_RNAi * k_on_RNAi - k_off_RNAi

k_on_miRNA <- 0.2e8 / (mole_unit_from_mole * time_unit_from_sec)
k_off_miRNA <- 0.051e-2 / time_unit_from_sec

### Combinations of data we use
volume_3T3 <- 3e-12 #4.4e-12 
total_RNA_3T3 <- 5e5

volume_mouse_oocyte <- 260e-12


one_step_degradation <- function(t, state, parameters) {
  with(as.list(c(state, parameters)), {
    d_target <- synthesis -k_on * enzyme * target + k_off * complex 
    d_enzyme <- -k_on * enzyme * target + k_off * complex + k_cat * complex
    d_complex <- k_on * enzyme * target - k_off * complex - k_cat * complex
    list(c(d_target, d_enzyme, d_complex))
  })
}

plot_ode_result <- function(out, title) {
  out_tidy <- out %>% as.data.frame() %>% 
    gather(type, concentration, -time) %>%
    mutate(time_h = time / (3600 * time_unit_from_sec)) 
  
  
  uncleaved_plot <- out_tidy %>% 
    filter(type != "enzyme") %>%
    group_by(time_h) %>%
    summarise(all_uncleaved = sum(concentration)) %>%
    (function(data) { 
      ggplot(data,aes(x = time_h, y = all_uncleaved)) + geom_line() +
        ggtitle(paste0(title," uncleaved RNAs"))  +
        ylim(-1e-5, max(data$all_uncleaved)) 
    })

  print(uncleaved_plot)
  
  auxiliary_plot <- 
    ggplot(out_tidy, aes(x = time_h, y = concentration, color = type, linetype = type)) + geom_line() +
    ggtitle(paste0(title," all reactants") ) + 
    ylim(-1e-5, max(out_tidy$concentration))


  print(auxiliary_plot)
}
```

We take \(k\_{on}\) and \(k\_{off}\) for the mouse miRNA system from from Wee, Flores-Jasso, Salomon & Zamore 2012, Argonaute Divides Its RNA Guide into Domains with Distinct Functions and RNA-Binding Properties, Figure 7. The same source also gives \(k\_{cat}\) for the RNAi pathway, which provides an upper bound on miRNA pathway efficiency.

We note that the \(k\_{cat}\) value for the RNAi pathway in the source (\(8.1\times 10^{-4} s^{-1}\)) is about a factor of 3.5 smaller from what would be calculated from the reported \(k\_{on}\), \(k\_{off}\) and \(K\_{M}\) (\(0.00283 s^{-1}\)). This is because \(k\_{on}\), \(k\_{off}\) were determined in a different experiment than \(K\_M\) and \(k\_{cat}\), but gives a useful insight into generalizability of the value.

For the following we use the RNAi \(k\_{cat}\) as reported.

```
miRNA_sim_single <- function(condition, times, init_complex_equilibrium = TRUE) {
  params <- c(k_cat = k_cat_RNAi * condition$k_cat_coeff, 
              k_on = k_on_miRNA, 
              k_off = k_off_miRNA,
              synthesis = condition$synthesis
              )
  
  total_target <- condition$total_target
  total_enzyme <- condition$total_enzyme
  
  
  if(init_complex_equilibrium) {
    #Init the enzyme and complex at equilibrium
    

    # Computing the initial complex concentration at equilibrium
    
    # K_D = k_on / k_off
    # c/t*e = K_D
    # c = t*e*K_D
    # 
    # t  = t_i - c
    # e  = e_i - c
    # 
    # c = (t_i - c)*(e_i-c)*K_D
    # 
    # c = c^2 * K_D + c*(-e_i -t_i)*K_D + e_i * t_i * K_D
    # 0 = c^2 * K_D + c*((-e_i -t_i)*K_D - 1) + e_i * t_i * K_D
    # 
    # D = ((-e_i -t_i)*K_D - 1)^2 - 4*e_i * t_i * K_D^2
    # c = ( -((-e_i -t_i)*K_D - 1)  +- sqrt(D) ) / (2 * K_D)
    # c = ( (e_i + t_i) * K_D + 1 + sqrt(D) ) / (2 * K_D)

    K_D <-  params["k_on"] / params["k_off"]
    names(K_D) <- NULL #For some reason, this has name and it messes up later computations
    complex_D <- (((total_target + total_enzyme) * K_D + 1) ^ 2) - 4 * total_target * total_enzyme * (K_D^2)
    initial_complex <- ((total_target + total_enzyme) * K_D + 1 - sqrt(complex_D)) / (2 * K_D)
    
    free_target <- total_target - initial_complex
    free_enzyme <- total_enzyme - initial_complex
    
    #Check that the equilibirum calculation is OK
    if(free_enzyme < 0 || free_target < 0 || initial_complex < 0) {
      stop("Concentrations below zero")
    }
    check <- initial_complex / (free_target * free_enzyme)
    rel_diff <- abs(check - K_D) / K_D
    if(rel_diff > 1e-3) {
      stop(paste("Not equilibrium, K_D =  ",K_D, "check = ", check, "  diff = ", rel_diff))
    }
  
    state <- c(target = free_target, enzyme = free_enzyme, complex = initial_complex)
  } else {
    initial_complex = 0
    state <- c(target = total_target, enzyme = total_enzyme, complex = 0)
  }
  
  out <- ode(y = state, times = times, func = one_step_degradation, parms = params)

  tidy_out <- out %>% as.data.frame() %>%
    gather(type, concentration, -time) %>%
    mutate(time_h = time / (3600 * time_unit_from_sec), initial_target = state["target"]) 

  crossing(as_tibble(condition), tidy_out)
}
```

### Comparison of concentrations - let7

Since the paper primarily studies the let7 miRNA, we start with a simulation exploring the possible dynamics of a miRNA roughly similar to let7 under various concentrations of the miRNA and its targets.

```
log_scale = TRUE
N_steps = 100

normal_miRNA_cpc_3T3 = 38000
normal_miRNA_cpc_oocyte = 20000

min_miRNA_cpc_oocyte =  normal_miRNA_cpc_oocyte / 2


if(log_scale) {
  max_miRNA_cpc_3T3 = normal_miRNA_cpc_3T3 * 10
} else {
  max_miRNA_cpc_3T3 = normal_miRNA_cpc_3T3 * 2
}

min_miRNA_concentration = (min_miRNA_cpc_oocyte / mole_unit) / volume_mouse_oocyte
max_miRNA_concentration = (max_miRNA_cpc_3T3 / mole_unit) / volume_3T3

min_target_concentration = min_miRNA_concentration
max_target_concentration = max_miRNA_concentration * 100

if(log_scale) {
  miRNA_concentrations = seq(log(min_miRNA_concentration), log(max_miRNA_concentration), length.out = N_steps) %>% exp()
  target_concentrations = seq(log(min_target_concentration), log(max_target_concentration), length.out = N_steps) %>% exp()
} else {
  miRNA_concentrations = seq(min_miRNA_concentration, max_miRNA_concentration, length.out = N_steps)
  target_concentrations = seq(min_target_concentration, max_target_concentration, length.out = N_steps)
}

times_concentration_hours = seq(0, 25, length.out = 101)
times_concentration = times_concentration_hours * 60 * 60 * time_unit_from_sec


k_cat_coeffs_concentration <- 1 / c(1, 4, 10)

all_conditions_concentration <- tibble(total_enzyme = miRNA_concentrations) %>%
  crossing(tibble(total_target = target_concentrations)) %>%
  crossing(k_cat_coeff = k_cat_coeffs_concentration) 
  
cell_info = tibble(condition = c("let7 in 3T3", "let7 in oocyte"),
                   cpc = c(normal_miRNA_cpc_3T3, normal_miRNA_cpc_oocyte),
                   volume = c(volume_3T3, volume_mouse_oocyte)) %>%
  mutate(miRNA_concentration = c(cpc / (mole_unit * volume))) %>% arrange(miRNA_concentration)
```

To determine which concentration ranges should we test, we take the measurements of copies per cell (cpc) of the let7 miRNA in 3T3 somatic cells and oocytes:

```
cell_info %>% mutate(`copies per cell` = cpc, `volume [pl]` = volume * 10^12, `concentration [nM]` = miRNA_concentration) %>% select(-volume, -cpc, -miRNA_concentration) %>% kable()
```

| condition | copies per cell | volume [pl] | concentration [nM] |
| --- | --- | --- | --- |
| let7 in oocyte | 20000 | 260 | 0.1277368 |
| let7 in 3T3 | 38000 | 3 | 21.0339865 |

This gives us anchor points to include in the simulations - we ended up testing miRNA concentrations in the range from 0.064nM to 0.2μM. We also do not know, how many let7 targets there are, so we explore range from very low target concentration (0.064nM) to much larger concentration of targets than let7 (21μM).

We assume the miRNA \(k\_{cat}\) to be lower than in RNAi, via relative efficiency koefficient \(r\) where \(k\_{cat,\mathrm{miRNA}} = r \cdot k\_{cat,\mathrm{RNAi}}\). Here we check \(r\) to be either 1, 0.25 or 0.1. We ran a simulation of the kinetics for each combination of those possible parameters.

```
#Note: this may taka quite a while, but not paralellized to avoid additional dependencies.

read_from_file <- FALSE
temp_file <- paste0(temp_dir, "/concentration.rds")
if(file.exists(temp_file)) {
  results_from_file <- readRDS(temp_file)
  if(nrow(results_from_file) == nrow(all_conditions_concentration) * length(times_concentration) * 3 && 
     identical(all_conditions_concentration, results_from_file %>% select(total_enzyme, total_target, k_cat_coeff) %>% distinct())) {
    read_from_file <- TRUE
    sim_results_concentration <- results_from_file
  } else {
    warning("Data read, but are different")
  }
}

if(!read_from_file) {
  sim_results_concentration <- all_conditions_concentration %>%
    mutate(synthesis = 0
           ) %>%
    rowwise() %>% do(miRNA_sim_single(., times_concentration, init_complex_equilibrium = TRUE)) %>% ungroup()
  
  saveRDS(sim_results_concentration, temp_file)
}
```

To see the effect we plot the proportion of the target mRNAs that is left uncleaved 20 hours relative to the initial concentration.

```
my_break_function <- function(limits) {
  min_order <- round(log10(limits[1]))
  max_order <- round(log10(limits[2]))
  10 ^ (min_order:max_order)
}

plot_efficiency_surface <- function(to_show, time_h, fixed_data, for_paper = FALSE) {
  if(log_scale) {
    scale_trans = "log10"
  } else {
    scale_trans = "identity"
  }

  
 if(for_paper) {
   fixed_data_highlight <- NULL
 } else {
    fixed_data_highlight <- geom_rect(data = fixed_data, mapping = aes(xmin = miRNA_concentration * 0.75, xmax = miRNA_concentration * 1.33, ymin = min_target_concentration / 1.5, ymax = max_target_concentration * 1.5), color = "lightgreen", fill = "transparent", inherit.aes = FALSE)
 }
  
 p <- to_show %>% filter(type == "target", time_h == !!time_h) %>% 
   mutate(concentration = if_else(concentration < 0, 0, concentration),
          relative_concentration = concentration / total_target) %>%
  ggplot(aes(x = total_enzyme, y = total_target, fill = relative_concentration, z = relative_concentration)) + 
   geom_raster() + 
   geom_point(data = fixed_data, mapping = aes(x = miRNA_concentration, y = min_target_concentration / 3, shape = condition), inherit.aes = FALSE, size = 2) +
   fixed_data_highlight +
   scale_fill_gradient("Target uncleaved", high = "#000000", low = "#f8f8f8", limits = c(0,1)) +
   scale_linetype_manual(values = c("solid","dotted")) + 
   scale_shape_manual("Condition", values = c("triangle","triangle open")) +
   facet_wrap(~k_cat_coeff, labeller= labeller(k_cat_coeff = function(x) {paste0("Rel. eff. = ",x)})) +
   scale_x_continuous("miRNA concentration [nM]", trans = scale_trans, breaks = my_break_function) +
   scale_y_continuous("Initial target concentration [nM]", trans = scale_trans, breaks = my_break_function) +
   ggtitle(paste0("Proportion target uncleaved at ", time_h,"h"))

 p  
}

plot_efficiency_surface(sim_results_concentration, 20, cell_info)
```

```
concentration_plot_for_paper <- plot_efficiency_surface(sim_results_concentration, 20, cell_info, for_paper = TRUE)

#Saving the plot for publication
for(type in c(".png",".svg")) {
  ggsave(paste0(temp_dir,"/simulation_concentrations", type), concentration_plot_for_paper, width = 9, height = 3.5)
}
```

In the plot above, the triangles and green rectangles highlight the area of the heatmaps that correspond to the expected let7 concentrations in 3T3 and oocytes. The main takeaway is that across the whole range of relative efficency, there is a big range of initial target concentrations where we would observe almost no repression in oocytes while the target would be almost completely repressed in 3T3 cells, simply due to stochiometry.

### Comparison of cell types

```
cells <- tibble(
  name = c("3T3","mouse oocyte"),
  volume = c(volume_3T3, volume_mouse_oocyte),
  total_RNA = c(total_RNA_3T3, 25e6),
  excess_RNA_synthesis = c(0, 0)
) %>% mutate(
  name = factor(name, levels = name)
)


target_pools <- tibble(fraction_of_total_RNA = c(0.1,0.2))

#Used with Shubha for experimental guess
#miRNA_counts <- tibble(miRNA_count = c(42000, 1e5, 5e5, 1e6, 5e6))
#miRNA_counts <- tibble(miRNA_count = c(2e4,5e4, 1e5, 1e6, 5e6))
miRNA_counts <- tibble(miRNA_count = c(2e4,5e4))

times_cells <- seq(0, 25 * 60 * 60 * time_unit_from_sec, length.out = 101)
time_step <- diff(times_cells)[1]


k_cat_coeffs <- tibble(k_cat_coeff = seq(0.05,1, length.out = 101))


all_conditions_cells <- crossing(cells, target_pools, miRNA_counts, k_cat_coeffs)
```

An additional way to look at the same results is to not vary the concentrations directly, but start with copies per cell in the individual cell types. Here, we will treat all the miRNA species as interchangeable. We test a range of relative efficiencies, starting at 0.05 and ending at 1.

We test dynamics in two cell types (3T3 and oocyte) as shown bellow::

```
cells %>% transmute(name = name, `total RNA [cpc]` = total_RNA, `volume [pl]` = volume * 10^12) %>%  kable()
```

| name | total RNA [cpc] | volume [pl] |
| --- | --- | --- |
| 3T3 | 5.0e+05 | 3 |
| mouse oocyte | 2.5e+07 | 260 |

We test different counts (cpc) of the miRNA species:

```
miRNA_counts %>% kable()
```

miRNA\_count

20000

50000

We test different amount of target mRNAs as fraction of the total RNA in the cell:

```
target_pools %>% kable()
```

fraction\_of\_total\_RNA

0.1

0.2

We solve the differential equation system for all possible combination of those values.

```
read_from_file <- FALSE
temp_file <- paste0(temp_dir, "/cells.rds")
if(file.exists(temp_file)) {
  results_from_file <- readRDS(temp_file)
  if(nrow(results_from_file) == nrow(all_conditions_cells) * length(times_cells) * 3 && 
     identical(all_conditions_cells, results_from_file %>% select(name, volume, total_RNA, excess_RNA_synthesis, fraction_of_total_RNA, miRNA_count, k_cat_coeff) %>% distinct())) {
    read_from_file <- TRUE
    sim_results_cells <- results_from_file
  } else {
    warning("Data read, but are different")
  }
}

if(!read_from_file) {
  sim_results_cells <- all_conditions_cells %>%
    mutate(synthesis = (excess_RNA_synthesis / mole_unit) * (fraction_of_total_RNA / volume),
           total_target = (total_RNA / mole_unit) * (fraction_of_total_RNA / volume),
           total_enzyme = miRNA_count / (volume * mole_unit)
  
           ) %>%
    rowwise() %>% do(miRNA_sim_single(., times_cells, init_complex_equilibrium = TRUE)) %>% ungroup()
  
  saveRDS(sim_results_cells, temp_file)
}
```

Below, we show the landscapes of concentrations of free miRNA targets over time for various relative efficiencies. Each heatmap corresponds to one of the simulated conditions (fraction of total RNA that are targets, miRNA count, type of cell). The concentration is reported relative to initial concentration of the free miRNA.

```
plot_concentration_surface <- function(to_show) {
  if(length(unique(to_show$fraction_of_total_RNA)) != 1) {
    stop("Has to be unique")
  }
  
  label_miRNA_count <- function(value) {
    value_proper = if_else(as.numeric(value) < 1e6, paste0(round(as.numeric(value)/1000),"K"),
      paste0(round(as.numeric(value)/1e6),"M"))
    
    paste0("miRNA cpc = ", value_proper)
  }
  
 p <- to_show %>% filter(type == "target") %>% group_by(fraction_of_total_RNA, miRNA_count, name) %>% 
   mutate(concentration = if_else(concentration < 0, 0, concentration),
          relative_concentration = concentration / max(concentration)) %>%
  ungroup() %>%
  ggplot(aes(x=time_h, y = k_cat_coeff, fill = relative_concentration, z = relative_concentration)) + 
   geom_raster() + 
   #scale_fill_distiller(palette = "Spectral") +
      scale_fill_gradient("Target uncleaved", low = "#505050", high = "#f0f0f0", limits = c(0,1)) +
   facet_grid(miRNA_count~name, labeller= labeller(miRNA_count = label_miRNA_count)) +
   scale_y_continuous("relative efficiency") +
   scale_x_continuous("time [h]") +
   ggtitle(paste("targets/total RNA =", to_show$fraction_of_total_RNA[1]))
  print(p)
  data.frame()
}

sim_results_cells %>%group_by(fraction_of_total_RNA) %>% do(plot_concentration_surface(.)) %>% invisible()
```

We see that even with the exact same miRNA degradation kinetics, all target mRNA is expected to be quickly degraded in 3T3 cells across a broad range of conditions, while in oocytes we don’t see noticeable degradation in most conditions.

### Conclusions

Both views of the simulations show that it is plausible for the differing stochiometry to account for differing miRNA activity in somatic cells and oocytes.

### Original computing environment

```
sessionInfo()
```

```
## R version 3.6.0 (2019-04-26)
## Platform: x86_64-w64-mingw32/x64 (64-bit)
## Running under: Windows 10 x64 (build 17134)
## 
## Matrix products: default
## 
## locale:
## [1] Slovak_Slovakia.1250
## 
## attached base packages:
## [1] stats     graphics  grDevices utils     datasets  methods   base     
## 
## other attached packages:
## [1] gdtools_0.1.8 knitr_1.23    cowplot_0.9.4 tibble_2.1.1  tidyr_0.8.3  
## [6] dplyr_0.8.1   ggplot2_3.1.1 deSolve_1.21 
## 
## loaded via a namespace (and not attached):
##  [1] Rcpp_1.0.1       pillar_1.4.0     compiler_3.6.0   plyr_1.8.4      
##  [5] highr_0.8        tools_3.6.0      digest_0.6.18    evaluate_0.13   
##  [9] gtable_0.3.0     pkgconfig_2.0.2  rlang_0.3.4      yaml_2.2.0      
## [13] xfun_0.7         withr_2.1.2      stringr_1.4.0    grid_3.6.0      
## [17] tidyselect_0.2.5 svglite_1.2.2    glue_1.3.1       R6_2.4.0        
## [21] rmarkdown_1.12   reshape2_1.4.3   purrr_0.3.2      magrittr_1.5    
## [25] scales_1.0.0     htmltools_0.3.6  assertthat_0.2.1 colorspace_1.4-1
## [29] labeling_0.3     stringi_1.4.3    lazyeval_0.2.2   munsell_0.5.0   
## [33] crayon_1.3.4
```
