## Supplementary code file for "Low miRNA abundance disables microRNA pathway in mammalian oocytes": README.md

# Supplementary code for "Low miRNA abundance disables microRNA pathway in mammalian oocytes"

- File "miRNA-sim.Rmd" contains full source code for the mathematical simulations used in the paper in the R markdown format.
- File "miRNA-sim.html" is the compiled report from "miRNA-sim.Rmd" showing all the results.
